# Circadian Rhythms in Murine Ocular Tissues including Sclera are affected by Neurobasal A Medium Preincubation, Mouse Strain, but not Sex

**DOI:** 10.1101/2024.03.01.583036

**Authors:** Nemanja Milićević, Cristina Sandu, Etienne Challet, Teemu O. Ihalainen, Soile Nymark, Marie-Paule Felder-Schmittbuhl

## Abstract

**Purpose:** Our understanding of ocular clocks has been profoundly advanced by the development of real-time recording of bioluminescence of PER2::LUC knock-in mouse explants. However, the effect of sex, mouse strain and culturing conditions on ocular clocks remains unknown. Here, we studied the role these variables play on PER2::LUC bioluminescence rhythms of ocular tissues: retinas, corneas and posterior eye cups (PEC). We also tested the hypothesis that the sclera contains a circadian oscillator by using scraped PEC as a proxy.

**Methods:** Retinas, corneas, intact and scraped PECs were obtained from male and female PER2::LUC knock-in mice maintained on either a pigmented C57BL/6J or albino RjOrl:SWISS background. PER2::LUC bioluminescence rhythms in ocular tissues were measured using a Lumicycle®.

**Results:** We compared PER2::LUC bioluminescence rhythms between ocular tissues and found that all ocular tissues oscillated, including the scraped PEC, which was previously not known to oscillate. The rhythms in scraped PECs had lower amplitudes, longer periods and distinct acrophases compared to other ocular tissues. Ocular tissues of RjOrl:SWISS mice oscillated with higher amplitudes compared to the ones of C57BL/6J, with corneal rhythms being most affected by mouse strain. A 24h preincubation with Neurobasal A medium enhanced rhythms of ocular tissues, whereas sex differences were not detected for these rhythms.

**Conclusions:** We discovered a novel oscillator in the sclera. PER2::LUC bioluminescence rhythms in murine ocular tissues are enhanced by Neurobasal A medium preincubation, mouse strain but not sex.

## Introduction

Rhythmic 24h changes in the Earth’s environment are one of the most salient selection pressures for all living beings. These challenges were met by the development of circadian rhythms i.e. innate, 24h biological cycles that govern the physiological, behavioral, and biochemical processes of living organisms^1^. In mammals, these rhythmic changes are orchestrated by the “central clock” located in suprachiasmatic nucleus (SCN) in the brain, which, in turn, coordinates the timing of peripheral clocks^2^. The core molecular components generating these oscillations are virtually the same in all cell types with transcription factors such as PER1-2, CLOCK, BMAL1, CRY1-2, REV-ERBs and RORs driving 24h periodicity of transcription-translation via negative interlocking feedback loops^3^. The coordinated activity of these factors drives rhythmic expression of “clock-controlled genes”, thereby enabling rhythmic adaptations in physiological, behavioral, and biochemical processes^3^.

The eye is considered as a peripheral clock with unique properties: It lies in direct contact with the principal time-giver (*Zeitgeber*) – light, and it is independent from the central clock, the SCN^4, 5^. Our understanding of ocular clocks has been profoundly advanced by the development of real-time monitoring of bioluminescence of PER2::LUC knock-in mouse explants ^6, 7^. Circadian rhythms were found in numerous ocular tissues, including the retina ^8-10^, cornea ^6, 11^, posterior eye cup (PEC) as a proxy for the retinal pigment epithelium (RPE) ^12^ and ciliary body ^13, 14^. Substantial effort was made in elucidating the regulation of these clocks. For example, it was found that the phase of ocular clocks is set by secreted neurohormones and neurotransmitters, such as dopamine as a day signal and melatonin as a night signal ^4, 13^. Each ocular clock is regulated in a distinct way, with dopamine being critical for entraining (i.e. setting the phase of) retinal ^9^ and PEC rhythms ^15^, whereas melatonin for corneal rhythms ^11, 16^.

There is considerable variability in culturing protocols of PER2::LUC explants, with some groups using a 24h preincubation with Neurobasal A medium and continued culturing using other media ^9, 17-19^, or without preincubation and culturing with M199 medium ^7^ or DMEM medium ^6, 16, 20^. Little is known about the influence these conditions play in PER2::LUC bioluminescence recordings. Our understanding of mouse strain effect on ocular clocks is also limited. Retinal explants of C57BL/6 and B6C3 mice show similar rhythms ^9^, whereas the regulation of corneal rhythms of C57BL/6 and C3Sn mice is distinct ^16^. To the best of our knowledge, there have been no studies on ocular clocks using mice with an albino background. Furthermore, the role of animal sex on ocular clocks has been largely ignored. Recent work showed no differences between male and female retinal rhythms ^21^. However, it is unknown whether sex affects corneal or PEC rhythms.

In this paper, we studied PER2::LUC bioluminescence rhythms in various ocular tissues: corneas, retinas, posterior eye cups (PEC) and scraped PECs as a proxy for sclerae. We then tested the possibility that PER2::LUC mice on albino RjOrl:SWISS background differ in bioluminescence rhythms in ocular tissues compared to pigmented C57BL/6J PER2::LUC mice. Finally, we explored the role culturing conditions and sex differences play in PER2::LUC bioluminescence rhythms in ocular tissues.

## Materials and Methods

### Mice

All experimental protocols were carried out according to the European Parliament and The Council of the European Union Directive (2010/63/EU) and institutional ethical guidelines (E-67-218-38). Animals were maintained in the Chronobiotron animal facility (UMS 3415, Strasbourg) under 12-h/12-h LD cycle (300 lx and <5 lx dim red light during light and dark, respectively) at a 22 ± 1 °C ambient temperature, with free access to food and water. Mice carrying the PER2::LUC reporter (Yoo et al., 2004) were bred either on a pigmented C57BL/6J (Charles River Laboratories, France) or albino RjOrl:SWISS (Janvier labs, Le Genest-Saint-Isle, France) background. We used both male and female mice. We used age matched 7-11 week old PER2::LUC mice maintained on C57BL/6J background. We also included 7-22 week old age-matched mice in experiments involving both C57BL/6J and RjOrl:SWISS backgrounds (**Fig. 3**). In **Figs. 1-3**, we used explants obtained from homozygous *mPer2*^*Luc*^ mice, whereas in Fig. 4, we obtained explants from heterozygous *mPer2*^*Luc*/+^ mice due to difficulties in breeding animals. Mice were sacrificed at varying times of day by cervical dislocation, eyes were enucleated and placed in HBSS media containing 4.2 mM sodium bicarbonate (Sigma S8761), 10 mM N-2-hydroxyethylpiperazine-N’-2-ethanesulfonic acid (Sigma H0887), 100 U/ml penicillin and 100 μg/ml streptomycin (Sigma P4333) as described previously ^17, 22, 23^.

**Figure 1.**
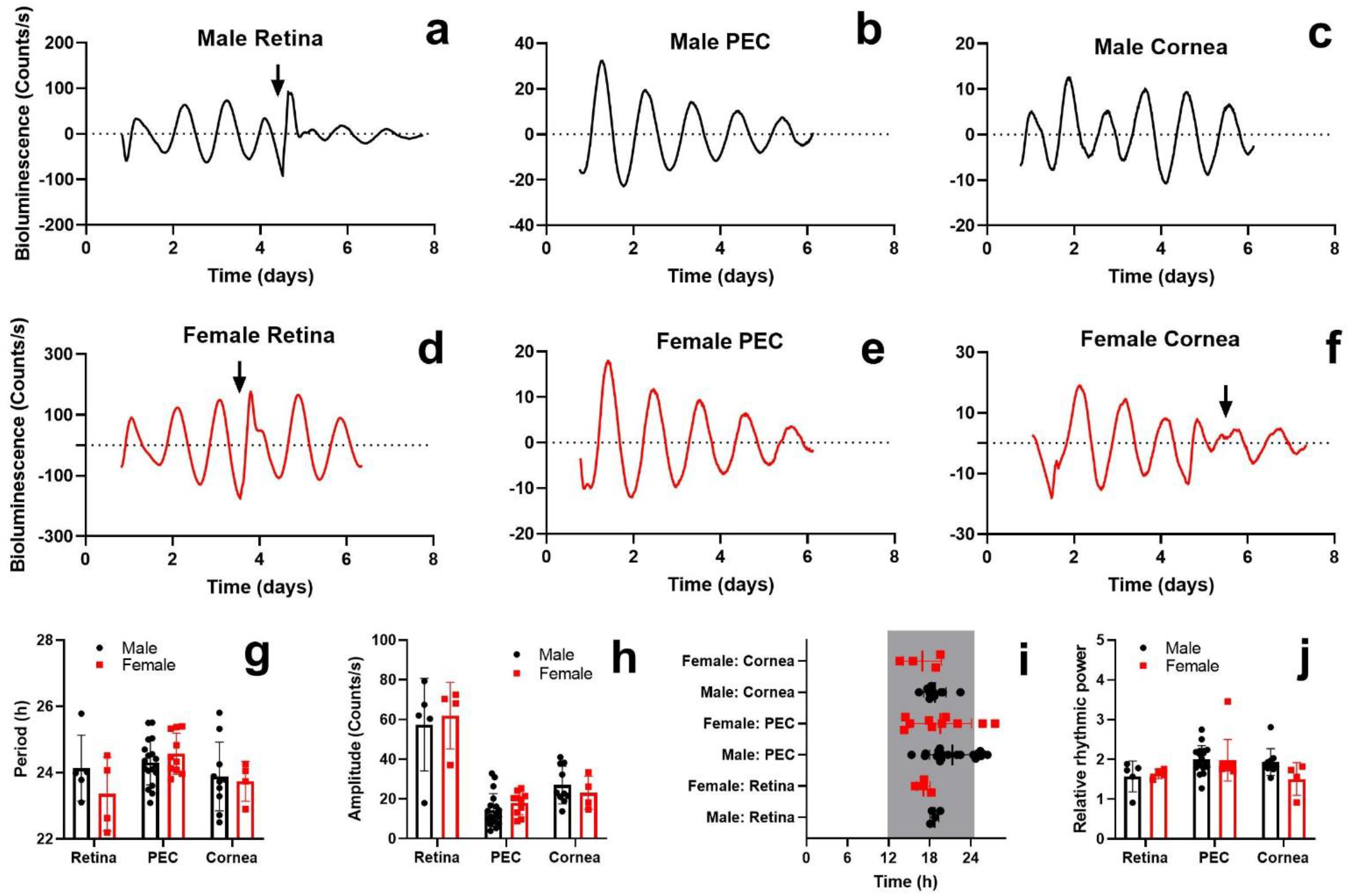
Sex does not affect circadian rhythms of PER2::LUC bioluminescence of *ex vivo* ocular tissues. Representative examples of bioluminescence rhythms in retina, PEC and cornea of (a - c) male and (d – f) female PER2::LUC mice bread on a C57BL/6J background. Animal sex did not significantly affect the following parameters of PER2::LUC bioluminescence rhythms: (g) period length, (h) amplitude, (i) acrophase, and (j) relative rhythmic power. The arrows indicate media change. Time in (i) is projected ZT with ZT12 = lights off. The gray rectangle represents subjective night-time. Individual data points are plotted together with means ± SD. PEC – posterior eye cup. N = 4 – 18

**Figure 2.**
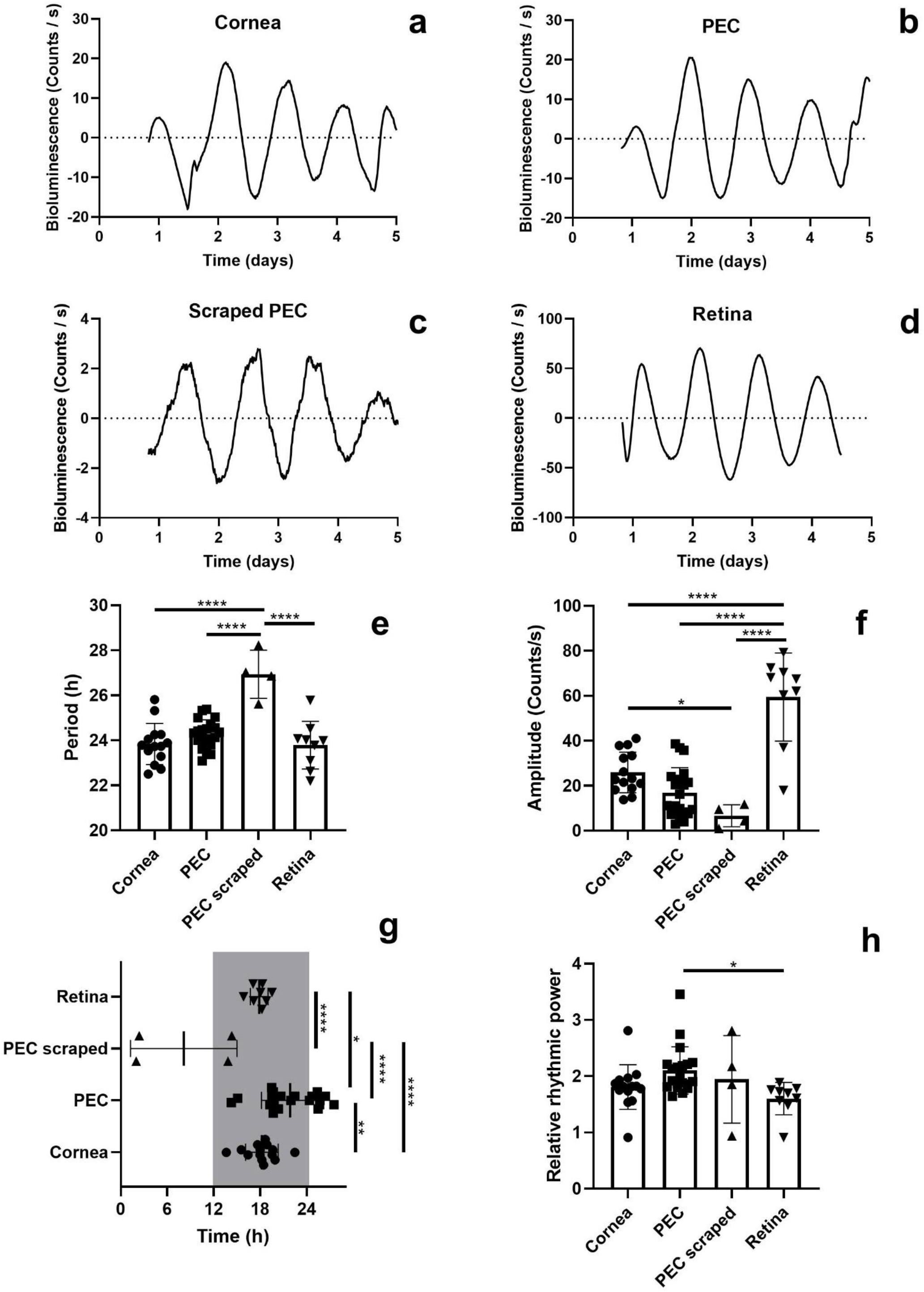
Circadian Rhythms of PER2::LUC Bioluminescence in *ex vivo* Ocular Tissues. Representative examples of PER2::LUC rhythms in (a) cornea, (b) intact PEC, (c) scraped PEC and (d) retina of mice bred on a C57BL/6J background. The bioluminescence rhythms of scraped PEC show (e) longer periods and (f) lower amplitudes compared to other ocular tissues. (g) The first obvious peak of PER2::LUC bioluminescence occurred earlier within the 24h cycle in scraped PEC compared to other ocular tissues. (h) The bioluminescence rhythms of PECs oscillated with higher relative rhythmic power compared to retinas. Time in (g) is projected ZT with ZT12 = lights off. The gray rectangle represents subjective night-time. Individual data points are plotted together with means ± SD. Holm-Sidak’s post-hoc test comparison, * P < 0.05, ** P < 0.01, **** P < 0.0001. PEC – posterior eye cup. N = 4 - 20

**Figure 3.**
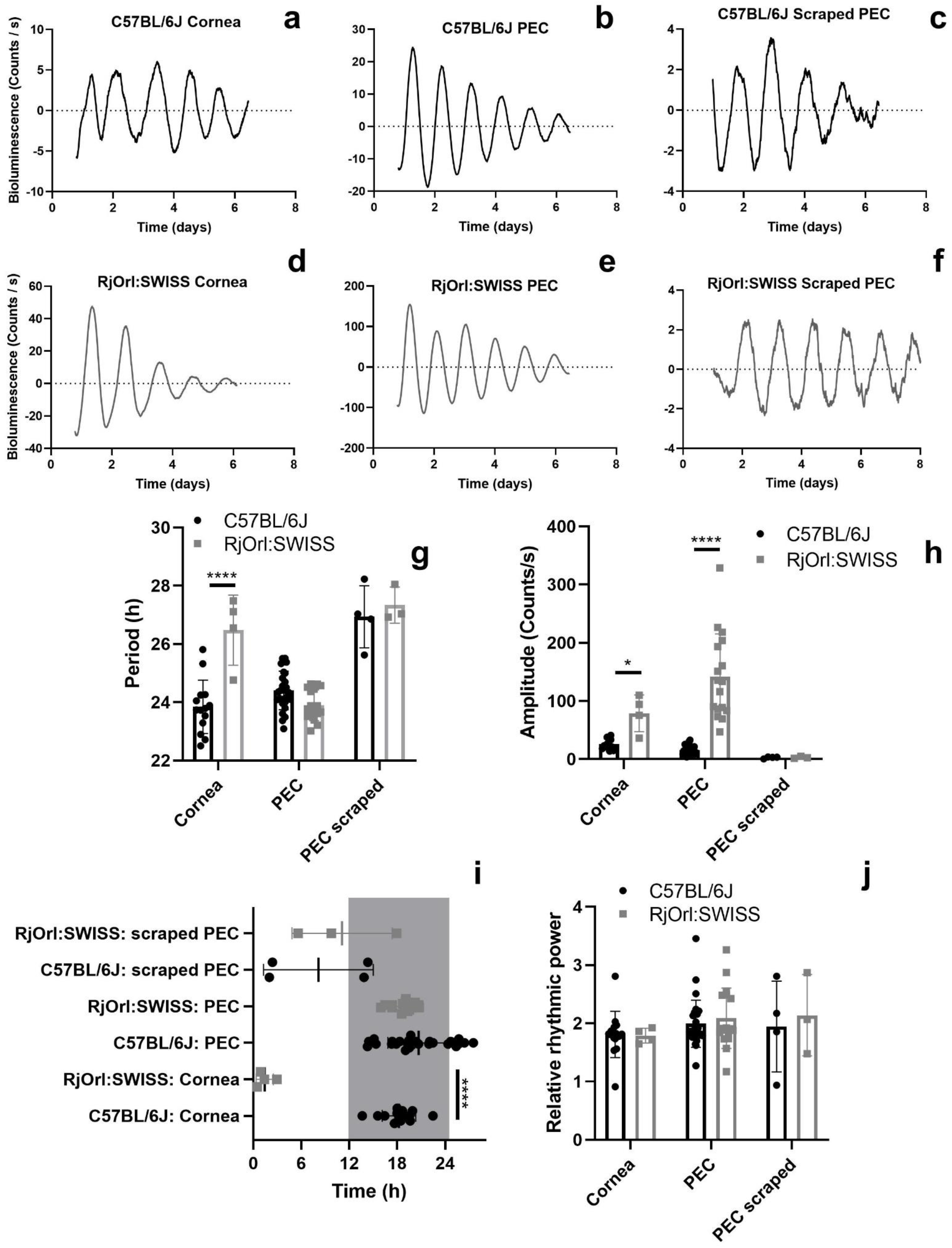
Mouse background influences circadian rhythms in ocular tissues. Representative examples of bioluminescence rhythms in corneas, intact PECs and scraped PECs obtained from PER2::LUC mice maintained on a (a-c) C57BL/6J background and, respectively, from a (d-f) RjOrl:SWISS background. Two-way ANOVA analysis was used to study the effects of background on (g) period length, (h) amplitude, (i) time of first peak i.e. acrophase and (j) relative rhythmic power of PER2::LUC bioluminescence oscillations in corneas, intact and scraped PECs. Time in (i) is projected ZT with ZT12 = lights off. The gray rectangle represents subjective night-time. Individual data points are plotted together with means ± SD. Holm-Sidak’s post-hoc test comparison, **** P < 0.0001. PEC – posterior eye cup. N = 3 – 28

**Figure 4.**
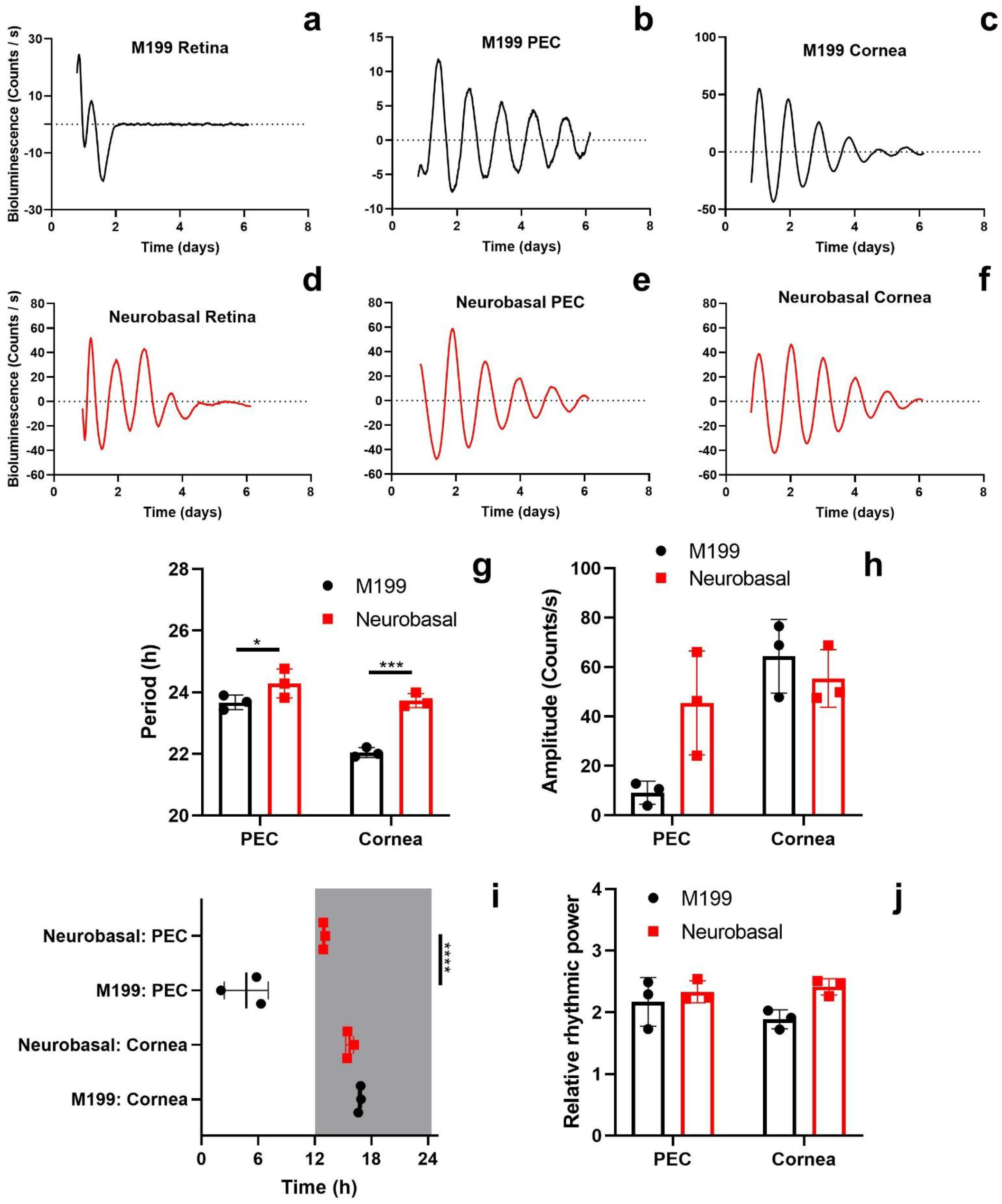
Preincubation medium has profound effects on circadian rhythms of PER2::LUC bioluminescence of *ex vivo* ocular tissues. Ocular tissues were obtained from heterozygous C57BL/6J *mPer2*^*Luc*/+^ mice. Representative examples of PER2::LUC bioluminescence rhythms in retinas, PECs and corneas that were preincubated for 24h with (a-c) M199 and (d-f) Neurobasal A medium. (g) Period length, (i) acrophase and (j) relative rhythmic power in ocular tissues was affected by the choice of preincubation medium. Conversely, (h) amplitude of PER2::LUC bioluminescence rhythms in ocular tissues was not affected by the choice of preincubation medium. Time in (i) is projected ZT with ZT12 = lights off. The gray rectangle represents subjective night-time. Individual data points are plotted together with means ± SD. Holm-Sidak’s post-hoc test comparison, * P < 0.05, **** P < 0.0001. PEC – posterior eye cup. N = 3

### Sample preparation

Dissections were performed in HBSS media at room temperature. A sterile needle was used to puncture the eyeball under the ora serrata and surgical scissors were used to cut around it. The lens and iris were dissected out of the cornea. The retina was excised from posterior eye cup. Removal of the RPE was performed by scraping of posterior eye cups by a curved blade and forceps. The retina, posterior eye cup and cornea were radially incised, flattened and placed on a semipermeable membrane (PICMORG50, EMD Millipore, Billerica, MA, USA) in a 35 mm culture dish.

### Bioluminescence recordings

Retinal samples were preincubated for 24 hours at 37°C in a humidified 5% CO_2_ atmosphere in 1 ml Neurobasal A medium (Invitrogen, Life Technologies, Carlsbad, CA, USA) containing antibiotics (25 U/ml penicillin and 25 mg/ml streptomycin), 2% B27 (Invitrogen, Life Technologies), and 2 mM L-glutamine. Then, the medium was replaced by 1 ml of medium 199 (Sigma-Aldrich, St. Louis, MO, USA) containing antibiotics (25 U/ml penicillin and 25 mg/ml streptomycin), 4 mM sodium bicarbonate, 20 mM D (+)-glucose, 2% B27, 0.7 mM L-glutamine, and 100 μM beetle luciferin (Promega, Fitchburg, WI, USA), and dishes were sealed (Dow Corning high-vacuum grease; Midland, MI, USA) under normal air and placed into the LumiCycle (Actimetrix, Wilmette, IL, USA) heated at 36°C. The procedure was modified for mouse posterior eye cups and corneas: preincubation for 24h and medium change was done in full 199 medium. All medium changes were performed under dim red light. Samples were recorded during at least 5 days: photons were counted during 112 seconds every 15 minutes.

Oscillations were analyzed using the Lumicycle Analysis software (Actimetrics, USA). Raw data (counts per second, cps) were baseline-subtracted (24-h running average, based on the whole recording period). LMFit (damped sine) function was used to calculate the periods and amplitudes with the following criteria for selecting the intervals: a minimum of least 3 cycles in length, up to 5 cycles, the first oscillation was excluded from the analysis and the selected interval had the maximum goodness of fit. The time of the first peak in the recordings is considered the acrophase and was extrapolated from the projected zeitgeber time (ZT), with ZT12 = lights off. The relative rhythmic power was determined using Periodogram function of Lumicycle Analysis to evaluate robustness of rhythms ^22, 24-27^.

### Statistics

Individual data points are plotted together with the means ± SD. Plots were generated using GraphPad Prism software (version 8.3.0, La Jolla, CA, USA). Normality of distribution was confirmed using the Kolmogorov-Smirnov test. One-way ANOVA analysis was performed in **Fig. 2**. Two-way ANOVA analyses were performed in this study with the following factors: **Fig. 1**: sex and tissue type; **Fig. 3**: mouse background and tissue type; **Fig. 4**: preincubation medium and tissue type. Further analyses, where indicated, were performed using the Holm-Sidak’s post-hoc tests. Student’s t-test was performed in **Fig S1**.

## Results

### Sex does not affect Circadian Rhythms of PER2::LUC Bioluminescence of *ex vivo* Ocular Tissues

We studied the effect of sex on ocular clocks in mice. We recorded PER2::LUC bioluminescence from C57BL/6J male retinas (**Fig. 1a**), PECs (**Fig. 1b**) and corneas (**Fig. 1c**) and female ones (**Figs 1d-f**). Retinal explants were preincubated with Neurobasal A medium, whereas PECs and corneas with M199 medium. Two-way ANOVA testing revealed that sex was not a statistically significant factor for period length (**Fig. 1g**; F(1, 45) = 0.70; P = 0.41), amplitude (**Fig. 1h**; F(1, 45) = 0.15; P = 0.70), acrophase (**Fig. 1i**; F(1, 44) = 2.78; P = 0.10) and relative rhythmic power (**Fig. 1j**; F(1, 45) = 1.03; P = 0.32) of PER2::LUC bioluminescence rhythms in C57BL/6J mice. We also studied PER2::LUC bioluminescence rhythms from male (**Fig. S1a**) and female (**Fig. S1b**) RjOrl:SWISS PECs. Similarly, there were no statistically significant differences in period length (**Fig. S1c**; Student’s t-test, t(15) = 0.23; P = 0.82), amplitude (**Fig. S1d**; Student’s t-test, t(15) = 0.85; P = 0.41), acrophase (**Fig. S1e**; Student’s t-test, t(15) = 1.88; P = 0.08) and relative rhythmic power (**Fig. 1Sf**; Student’s t-test, t(15) = 0.47; P = 0.65) between male and female PECs of RjOrl:SWISS mice. Overall, these results suggest that sex does not influence ocular clocks in mice. Bearing this conclusion in mind, we included results obtained from samples of both sexes in further experiments.

### Circadian Rhythms of PER2::LUC Bioluminescence in *ex vivo* Ocular Tissues: a Clock in the Sclera?

We initially recorded PER2::LUC bioluminescence from corneas (**Fig. 2a**), PECs (**Fig. 2b**) and retinas (**Fig. 2d**) obtained from PER2::LUC mice on the C57BL/6J background. It was implied that scraped posterior eye cups do not have measurable rhythms in bioluminescence^12^. Thus, we included scraped PECs as negative controls. Unexpectedly, we observed that sclerae showed circadian oscillations of PER2::LUC bioluminescence (**Fig. 2c**). We compared the rhythm in the sclera to oscillators of other ocular tissues: the cornea, posterior eye cup and retina. To increase statistical power, we pooled data from 7 experiments with ocular tissues being obtained from a total of 25 mice. One-way ANOVA analyses revealed that period length (**Fig. 2e**; F(3, 43) = 15.37; P < 0.0001), amplitude (**Fig. 2f**; F(3, 43) = 28.90; P < 0.0001), acrophase (**Fig. 2g**; F(3, 42) = 19.53; P < 0.0001) and relative rhythmic power (**Fig. 2h**; F(3, 43) = 3.14; P = 0.035) differed significantly between bioluminescence rhythms of different ocular tissues. Holm-Sidak’s post-hoc tests showed that scraped PECs had significantly longer periods compared to corneas (P < 0.0001), intact PECs (P < 0.0001) and retinas (P < 0.0001). In addition, the bioluminescence rhythms of scraped PECs had significantly lower amplitudes compared to the ones of corneas (P = 0.025) and retinas (P < 0.0001). The acrophases of scraped PECs rhythms differed significantly compared to corneas (P < 0.0001), intact PECs (P < 0.0001) and retinas (P < 0.0001). These results suggest that the sclera contains a circadian oscillator with lower amplitudes, longer periods and distinct acrophases compared to other ocular tissues.

### Does Mouse Strain affect Circadian Rhythms of PER2::LUC Bioluminescence of *ex vivo* Ocular Tissues?

We speculated that mouse background may influence ocular rhythms. To test this hypothesis, we measured bioluminescence rhythms from corneas (**Fig. 3a, d**), PECs (**Fig. 3b, e**) and scraped PECs (**Fig. 3c, f**) from PER2::LUC mice maintained on two backgrounds: the pigmented one, C57BL/6J, and the albino line RjOrl:SWISS. Two-way ANOVA testing revealed that mouse background was a significant factor for period length (**Fig. 3g**; F(1, 64) = 11.50; P = 0.0012), amplitude (**Fig. 3h**; F(1, 64) = 10.88; P < 0.0001), acrophase (**Fig. 3i**; F(1, 64) = 22.19; P < 0.0001), but not relative rhythmic power (**Fig. 3j**; F(1, 64) = 0.34; P = 0.33) of PER2::LUC bioluminescence rhythms. In particular, Holm-Sidak’s post-hoc testing revealed that corneas of RjOrl:SWISS mice have significantly longer periods compared to corneas of C57BL/6J mice (**Fig. 3g**; P < 0.0001). Furthermore, PECs (**Fig. 3h**; P < 0.0001) and corneas (**Fig. 3h**; P = 0.033) of RjOrl:SWISS mice have significantly higher amplitudes compared to the ones of C57BL/6J mice. Holm-Sidak’s post-hoc testing revealed that corneas of RjOrl:SWISS mice have significantly different acrophases compared to corneas of C57BL/6J mice (**Fig. 3i**; P < 0.0001). These results show that mouse background significantly affects PER2::LUC bioluminescence rhythms of ocular tissues. Among the studied ocular tissues, the rhythms in corneas were most affected by mouse background.

### Neurobasal A Medium preincubation enhances Circadian Rhythms of PER2::LUC Bioluminescence of *ex vivo* Ocular Tissues

A 24h preincubation with Neurobasal A medium is used for maintaining retinal explant viability for PER2::LUC bioluminescence recordings ^9^. Such a procedure is not necessary for studying rhythms in PECs ^12, 15, 23, 28^ and corneas ^11, 28^. We tested the hypothesis that Neurobasal A medium preincubation influences PER2::LUC bioluminescence rhythms in ocular tissues. Retinas, PECs and corneas were obtained from heterozygous C57BL/6J *mPer2*^*Luc*/+^ mice. These tissues were preincubated for 24h with either recording medium, M199 (representative PER2::LUC bioluminescence recordings of retina, PEC and cornea: **Figs 4a-c**) or Neurobasal A (retina, PEC and cornea: **Figs 4d-f**), followed by a medium change in all samples with recording medium containing M199. At least 3 complete cycles were observed in all bioluminescence recordings, except in the ones recorded from M199 preincubated retinas whose oscillations were highly variable and attenuated rapidly (**Fig. 4a**). Due to this reason, we could not accurately measure parameters of oscillations in M199 preincubated retinas. Thus, it was not possible to statistically compare Neurobasal A and M199 preincubated retinas. Conversely, such comparisons were possible for PEC and cornea recordings. Two-way ANOVA testing revealed that the choice of preincubation medium was a significant factor for period length (**Fig. 4g**; F(1, 8) = 44.82; P = 0.0002), acrophase (**Fig. 4i**; F(1, 8) = 27.01; P = 0.0008) and relative rhythmic power of PER2::LUC bioluminescence rhythms (**Fig. 4j**; F(1, 8) = 6.19; P = 0.038), but not amplitude (**Fig. 4h**; F(1, 8) = 2.72; P = 0.14). Holm-Sidak’s post-hoc testing revealed that PECs preincubated with Neurobasal A have significantly longer periods (P = 0.036) and distinct acrophases (P < 0.0001) compared to PECs preincubated with M199 medium. Although the two-way ANOVA analysis did not show an effect of preincubation medium on amplitudes, Holm-Sidak’s post-hoc test showed higher amplitudes (P = 0.029) of Neurobasal A preincubated PECs compared to M199 treated ones. We also found that corneas preincubated with Neurobasal A medium have significantly longer periods compared to the ones preincubated with M199 medium (P = 0.0002). These results show that Neurobasal A medium can enhance circadian rhythmicity not only in retinas, but also in other ocular tissues such as PECs and corneas.

## Discussion

In this study, we characterized PER2::LUC bioluminescence rhythms in various ocular tissues: corneas, retinas, PECs and discovered a hitherto unknown oscillator in the sclera. We did not observe any significant effect of sex in PER2::LUC expression in ocular tissues. Further, we showed the significance that *mPer2*^*Luc*^ mouse background plays in bioluminescence rhythms of ocular clocks. Finally, we found that culturing conditions, such as Neurobasal A medium preincubation, can substantially enhance rhythms of PER2::LUC bioluminescence of most ocular tissues.

Several lines of evidence suggested that ocular rhythms may differ between sexes. The diurnal variation of intraocular pressure and tear production was shown to vary between sexes^29, 30^. Androgen, estrogen and progesterone receptor mRNA was found in a variety of ocular tissues^31, 32^. In certain tissue types such as livers and adrenal glands, there were sex differences in PER2::LUC rhythms^33^. However, we did not find any significant sex differences in PER2::LUC bioluminescence rhythms of retinal, corneal and PEC explants. Calligaro and colleagues also showed no effect of sex on retinal rhythms^21^. Although our results show that, between sexes, there are no intrinsic differences in ocular rhythms, we cannot exclude the possibility that sex hormones may modulate these rhythms.

Prior work on the regulation of the RPE clock was conducted using bioluminescence recordings of PEC explants from *Per2*^*Luc*^ mice ^12, 15, 23, 28^. It was implied that scraping the RPE cell layer from the PEC attenuates oscillations in the explant, indicating that the RPE is the source of rhythms of PER2::LUC bioluminescence in the PEC^12^. Unexpectedly, we found that scraped PECs *did* show oscillations of PER2::LUC bioluminescence, suggesting that the sclera also contains a clock. These oscillations had lower amplitudes, longer periods and distinct acrophases compared to other ocular tissues, suggesting that this is a weak and less coupled oscillator^34^ compared to other ocular clocks. We do not know the cell types which are generating these rhythms. The sclera is comprised of collagen-rich scleral extracellular matrix and sparsely populated resident cells, called fibrocytes^35^. Confirming the presence of clock proteins with immunolabelling is complicated by the lack of specific markers for fibrocytes and their sparse localization^36^. However, it is known that, upon insult, fibrocytes can undergo transformation into active fibroblasts^35^, a cell type known to show robust circadian oscillations^37^. Therefore, it is likely that scleral rhythms are generated by resident fibrocytes.

We found that bioluminescence of corneas, retinas and PECs oscillated with a periodicity of roughly 24h, which is consistent with previous work ^28^. Also, the acrophase of retinal rhythms occurred roughly in the late night, as reported by others^16, 28^. Conversely, the peak timing of PEC bioluminescence rhythms is not consistent with literature^28^, whereas the acrophase of corneal rhythms varies depending on the study^16, 28^. The reason for such variability might be due to high dispersion of our data and/or the methods for calculating the acrophase. However, we think it is most likely due to differences in culturing protocols. In support of this explanation, we found that Neurobasal A medium preincubation can substantially influence PER2::LUC bioluminescence rhythms in ocular tissues. This protocol is widely used to enable long-term culture of retinal explants for monitoring PER2::LUC expression^9, 18, 21, 27, 38^, which was also essential for our prior work^17, 18, 22, 24^. Our results showed that Neurobasal A medium preincubated retinas exhibited consistent and robust oscillations. Neurobasal A medium preincubation increased the period lengths and altered the acrophases of corneal and PEC rhythms compared to the ones preincubated with M199. However, it was previously reported that recording PER2::LUC bioluminescence rhythms in retinal explants was possible with only M199 medium^7, 10, 28^. We found that bioluminescence rhythms in retinas preincubated with M199 medium gave highly variable rhythms which quickly attenuated. Conversely, in **Figs 1, 2** and **3**, the PEC and cornea explants still showed reasonably stable and consistent oscillations despite being cultured in M199 medium, as recommended by Baba and colleagues^7, 11, 12^. These effects could be explained by the fact that Neurobasal A medium contains a concentration of components optimized for culturing neurons^39^.

Most work using *Per2*^*Luc*^ mice to study ocular clocks was done on a C57BL6 backround^9-12, 15, 17, 19, 20, 22-24, 27, 28, 40^. There is a limited number of studies in which a different mouse strain was used to study ocular clocks^9, 16^. These studies have used such backgrounds as a strategy to overcome the main limitation of the C57BL6 in chronobiology, i.e. the inability of these mice to secrete the clock regulatory hormone, melatonin^41^. Recent work suggests that the effect of background on ocular clock function can be unrelated to melatonin deficiency. Specifically, C57BL/6 and 129T2/SvEmsJ mice are both deficient in melatonin^41-43^, and they showed differences in timing of phagocytosis of photoreceptor outer segments^44-46^, a known circadian clock-regulated process in the eye^47^. One such non-melatonin strain related feature which is associated with altered circadian rhythms in mice is pigment mutation^48^. In this manuscript, we studied ocular clocks using the albino RjOrl:SWISS background^49^, a line which has a similar free-running period as C57BL/6^50^. Our results showed significant differences in bioluminescence rhythms in ocular tissues of albino RjOrl:SWISS compared to pigmented C57BL/6J *mPer2*^*Luc*^ mice. RjOrl:SWISS corneas and PEC had higher amplitudes of PER2::LUC bioluminescence compared to the ones of C57BL/6J mice. Corneal bioluminescence of RjOrl:SWISS oscillated with a different phase and longer period compared to the ones of C57BL/6J mice suggesting that corneal clocks are more affected by pigment mutation. These differences may be due to technical or biological reasons. It is possible that the pigment of C57BL/6J ocular tissues may confound the PER2 bioluminescence signal, leading to lower amplitudes in these explants as observed in our results. This hypothesis does not explain the differences in corneal rhythms because the cornea is transparent^51^. To the best of our knowledge, the ability of RjOrl:SWISS to synthesize melatonin is not known. Melatonin secretion may explain the differences in corneal clocks between the strains, as the corneal clock is set by this hormone ^11, 16^.

There are limitations in our study. Our strategy for calculating acrophases is likely affected by period length. We also observed high variability in estimating the acrophase in scleral samples (**Fig. 2g** and **3i**). Despite our best efforts, we cannot fully exclude the possibility that these samples may contain remnants of RPE or choroid. The outliers have similar acrophases as intact PECs, which supports this possibility. By contrast, period lengths and amplitudes remained consistent in scleral rhythms, suggesting that they originate from the same cell type. We also cannot exclude the possibility that the scraping procedure could have affected PER2::LUC bioluminescence rhythms. A further limitation is the relatively wide age range of mice used in this study (7-22 weeks). It is reported that there are limited differences in PER2::LUC bioluminescence rhythms of retinas of mice aged 2-3 months compared to the ones aged up to 8 months^21^. Thus, it is unlikely that age differences confounded our results because we used age matched mice with the youngest ones being ∼2 months old. Next, we used homozygous *mPer2*^*Luc*^ mice for experiments shown in **Figs 1-3**, whereas heterozygous *mPer2*^*Luc*/+^ mice in **Fig. 4** due to difficulties in breading homozygous mice. These allele differences may explain the observed discrepancies in acrophases of ocular tissues in **Figs 2, 3** compared to **Fig 4**, slightly earlier periods in **Fig. 4** compared to **Figs 2, 3**; discrepancies in amplitude in **Fig. 4** compared to **Figs 1-3**. However, we used the experiments in **Fig. 4** to study the effects of culturing conditions on PER2::LUC bioluminescence. Thus, the observed discrepancies did not affect our conclusions.

This work opens the possibility for various future directions. The identification of cell types generating corneal rhythms, the regulation of this oscillator and the physiological functions which are under its control remain open questions. The component in Neurobasal A medium that enhances ocular rhythms remains elusive. We do not know the molecular mechanisms underlying the differences in ocular rhythms between mouse strains. These, along with other questions remain topics for future investigation.

## Supplemental material

**Figure S1.**
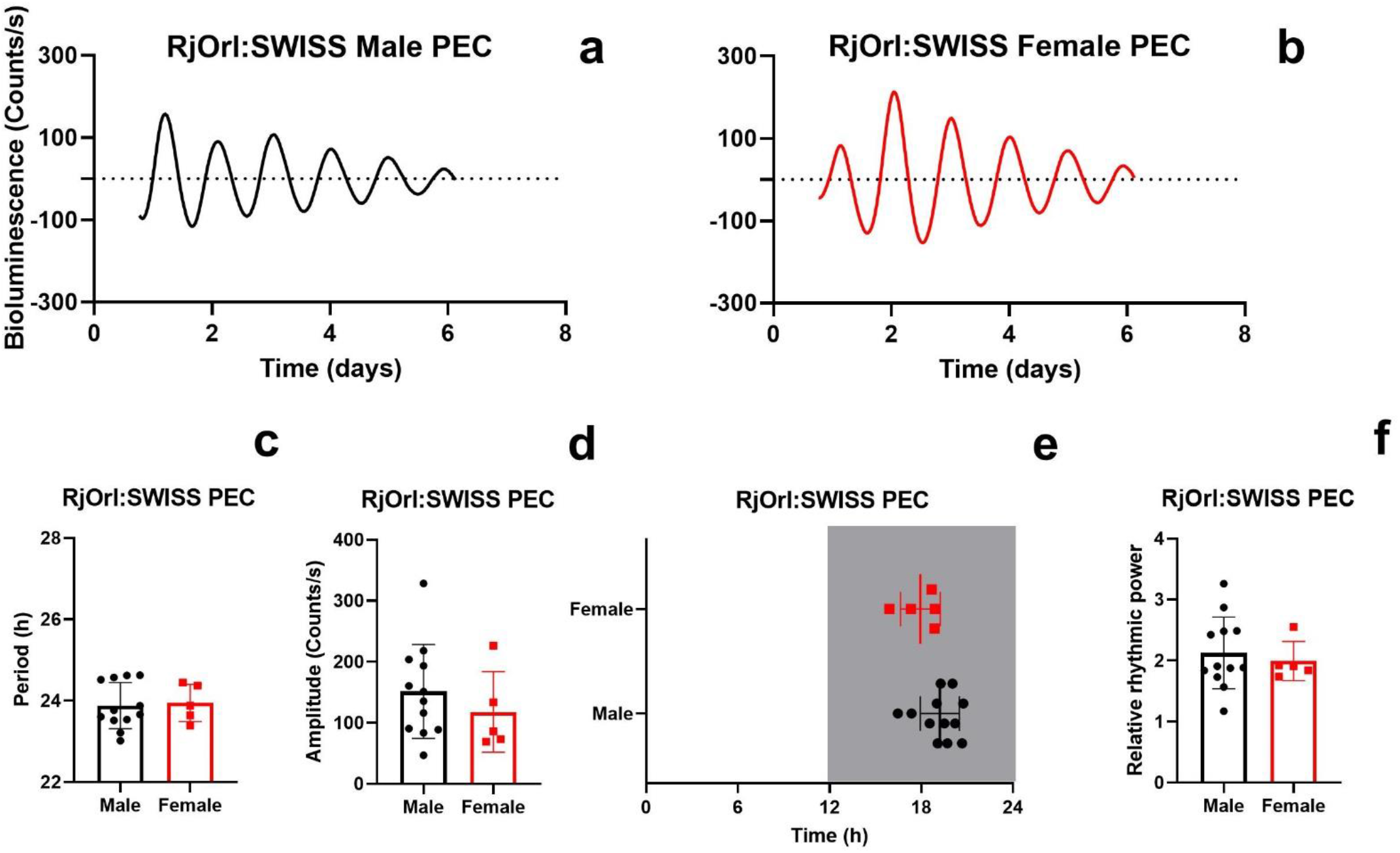
Sex does not affect circadian rhythms of PER2::LUC bioluminescence of PEC obtained from RjOrl:SWISS mice. Representative traces are shown of (a) male and (b) female PER2::LUC bioluminescence of PECs from RjOrl:SWISS mice. Animal sex did not significantly affect the (c) period length, (d) amplitude, (e) acrophase and (f) relative rhythmic power of PER2::LUC bioluminescence rhythms. Time in (e) is projected ZT with ZT12 = lights off. The gray rectangle represents subjective night-time. Individual data points are plotted together with means ± SD. PEC – posterior eye cup. N = 5 – 12

## Acknowledgments

This study was supported by the Centre National pour la Recherche Scientifique and the University of Strasbourg (France), the Finnish Cultural Foundation (Finland), the Federation of European Biochemical Societies (FEBS) Short-Term Fellowship (EU) and the Academy of Finland (decision number: 340127). We extend our gratitude to Dr D. Ciocca-Sage, Dr S. Reibel and N. Lethenet from the Chronobiotron (UMS 3415) for animal care.

## Data availability statement

Raw data that support the findings of this study are available from the corresponding authors, upon reasonable request.

## References

1. Cox KH, Takahashi JS. Circadian clock genes and the transcriptional architecture of the clock mechanism. Journal of molecular endocrinology 2019;63:R93–r102.

2. Cox KH, Takahashi JS. Introduction to the Clock System. Advances in experimental medicine and biology 2021;1344:3–20.

3. Takahashi JS. Molecular components of the circadian clock in mammals. Diabetes, obesity & metabolism 2015;17 Suppl 1:6–11.

4. McMahon DG, Iuvone PM, Tosini G. Circadian organization of the mammalian retina: from gene regulation to physiology and diseases. Progress in retinal and eye research 2014;39:58–76.

5. Felder-Schmittbuhl MP, Buhr ED, Dkhissi-Benyahya O, et al. Ocular Clocks: Adapting Mechanisms for Eye Functions and Health. Investigative ophthalmology & visual science 2018;59:4856–4870.

6. Yoo SH, Yamazaki S, Lowrey PL, et al. PERIOD2::LUCIFERASE real-time reporting of circadian dynamics reveals persistent circadian oscillations in mouse peripheral tissues. Proceedings of the National Academy of Sciences of the United States of America 2004;101:5339–5346.

7. Baba K, Tosini G. Real-Time Monitoring of Circadian Rhythms in the Eye. Methods in molecular biology (Clifton, NJ) 2022;2550:367–375.

8. Tosini G, Menaker M. Circadian rhythms in cultured mammalian retina. Science (New York, NY) 1996;272:419–421.

9. Ruan GX, Allen GC, Yamazaki S, McMahon DG. An autonomous circadian clock in the inner mouse retina regulated by dopamine and GABA. PLoS biology 2008;6:e249.

10. Ruan GX, Zhang DQ, Zhou T, Yamazaki S, McMahon DG. Circadian organization of the mammalian retina. Proceedings of the National Academy of Sciences of the United States of America 2006;103:9703–9708.

11. Baba K, Davidson AJ, Tosini G. Melatonin Entrains PER2::LUC Bioluminescence Circadian Rhythm in the Mouse Cornea. Investigative ophthalmology & visual science 2015;56:4753–4758.

12. Baba K, Sengupta A, Tosini M, Contreras-Alcantara S, Tosini G. Circadian regulation of the PERIOD 2::LUCIFERASE bioluminescence rhythm in the mouse retinal pigment epithelium-choroid. Molecular vision 2010;16:2605–2611.

13. Besharse JC, McMahon DG. The Retina and Other Light-sensitive Ocular Clocks. Journal of biological rhythms 2016;31:223–243.

14. Tsuchiya S, Buhr ED, Higashide T, Sugiyama K, Van Gelder RN. Light entrainment of the murine intraocular pressure circadian rhythm utilizes non-local mechanisms. PloS one 2017;12:e0184790.

15. Baba K, DeBruyne JP, Tosini G. Dopamine 2 Receptor Activation Entrains Circadian Clocks in Mouse Retinal Pigment Epithelium. Scientific reports 2017;7:5103.

16. Huynh AV, Buhr ED. Melatonin Adjusts the Phase of Mouse Circadian Clocks in the Cornea Both Ex Vivo and In Vivo. Journal of biological rhythms 2021;36:470–482.

17. Jaeger C, Sandu C, Malan A, Mellac K, Hicks D, Felder-Schmittbuhl MP. Circadian organization of the rodent retina involves strongly coupled, layer-specific oscillators. FASEB journal : official publication of the Federation of American Societies for Experimental Biology 2015;29:1493–1504.

18. Calligaro H, Coutanson C, Najjar RP, et al. Rods contribute to the light-induced phase shift of the retinal clock in mammals. PLoS biology 2019;17:e2006211.

19. Buhr ED, Yue WW, Ren X, et al. Neuropsin (OPN5)-mediated photoentrainment of local circadian oscillators in mammalian retina and cornea. Proceedings of the National Academy of Sciences of the United States of America 2015;112:13093–13098.

20. Díaz NM, Lang RA, Van Gelder RN, Buhr ED. Wounding Induces Facultative Opn5-Dependent Circadian Photoreception in the Murine Cornea. Investigative ophthalmology & visual science 2020;61:37.

21. Calligaro H, Kinane C, Bennis M, Coutanson C, Dkhissi-Benyahya O. A standardized method to assess the endogenous activity and the light-response of the retinal clock in mammals. Molecular vision 2020;26:106–116.

22. Gegnaw ST, Sandu C, Mendoza J, Bergen AA, Felder-Schmittbuhl MP. Dark-adapted light response in mice is regulated by a circadian clock located in rod photoreceptors. Experimental eye research 2021;213:108807.

23. Milicevic N, Mazzaro N, de Bruin I, et al. Rev-Erbalpha and Photoreceptor Outer Segments modulate the Circadian Clock in Retinal Pigment Epithelial Cells. Scientific reports 2019;9:11790.

24. Gegnaw ST, Sandu C, Mazzaro N, Mendoza J, Bergen AA, Felder-Schmittbuhl MP. Enhanced Robustness of the Mouse Retinal Circadian Clock Upon Inherited Retina Degeneration. Journal of biological rhythms 2022;37:567–574.

25. Buonfiglio DC, Malan A, Sandu C, et al. Rat retina shows robust circadian expression of clock and clock output genes in explant culture. Molecular vision 2014;20:742–752.

26. Herzog ED, Kiss IZ, Mazuski C. Measuring synchrony in the mammalian central circadian circuit. Methods in enzymology 2015;552:3–22.

27. Ruan GX, Gamble KL, Risner ML, Young LA, McMahon DG. Divergent roles of clock genes in retinal and suprachiasmatic nucleus circadian oscillators. PloS one 2012;7:e38985.

28. Baba K, Tosini G. Aging Alters Circadian Rhythms in the Mouse Eye. Journal of biological rhythms 2018;33:441–445.

29. Kulualp K, Kilic S, Aytekin O. Effects of sex, eye-side, diurnal variation on intraocular pressure in calves. Polish journal of veterinary sciences 2019;22:67–74.

30. Piccione G, Giannetto C, Fazio F, Giudice E. Daily rhythm of tear production in normal horse. Veterinary ophthalmology 2008;11 Suppl 1:57–60.

31. Wickham LA, Gao J, Toda I, Rocha EM, Ono M, Sullivan DA. Identification of androgen, estrogen and progesterone receptor mRNAs in the eye. Acta ophthalmologica Scandinavica 2000;78:146–153.

32. Kobayashi K, Kobayashi H, Ueda M, Honda Y. Estrogen receptor expression in bovine and rat retinas. Investigative ophthalmology & visual science 1998;39:2105–2110.

33. Kuljis DA, Loh DH, Truong D, et al. Gonadal- and sex-chromosome-dependent sex differences in the circadian system. Endocrinology 2013;154:1501–1512.

34. Schmal C, Herzog ED, Herzel H. Measuring Relative Coupling Strength in Circadian Systems. Journal of biological rhythms 2018;33:84–98.

35. Boote C, Sigal IA, Grytz R, Hua Y, Nguyen TD, Girard MJA. Scleral structure and biomechanics. Progress in retinal and eye research 2020;74:100773.

36. Reinhardt JW, Breuer CK. Fibrocytes: A Critical Review and Practical Guide. Frontiers in immunology 2021;12:784401.

37. Balsalobre A, Damiola F, Schibler U. A serum shock induces circadian gene expression in mammalian tissue culture cells. Cell 1998;93:929–937.

38. Kinane C, Calligaro H, Jandot A, et al. Dopamine modulates the retinal clock through melanopsin-dependent regulation of cholinergic waves during development. BMC biology 2023;21:146.

39. Brewer GJ, Torricelli JR, Evege EK, Price PJ. Optimized survival of hippocampal neurons in B27-supplemented Neurobasal, a new serum-free medium combination. Journal of neuroscience research 1993;35:567–576.

40. Goyal V, DeVera C, Baba K, et al. Photoreceptor Degeneration in Homozygous Male Per2(luc) Mice During Aging. Journal of biological rhythms 2021;36:137–145.

41. Ebihara S, Marks T, Hudson DJ, Menaker M. Genetic control of melatonin synthesis in the pineal gland of the mouse. Science (New York, NY) 1986;231:491–493.

42. Kasahara T, Abe K, Mekada K, Yoshiki A, Kato T. Genetic variation of melatonin productivity in laboratory mice under domestication. Proceedings of the National Academy of Sciences of the United States of America 2010;107:6412–6417.

43. Balazs I, Purrello M, Rocchi M, Rinaldi A, Siniscalco M. Is the gene for steroid sulfatase X-linked in man? An appraisal of data from humans, mice, and their hybrids. Cytogenet Cell Genet 1982;32:251.

44. Milićević N, Ait-Hmyed Hakkari O, Bagchi U, et al. Core circadian clock genes Per1 and Per2 regulate the rhythm in photoreceptor outer segment phagocytosis. FASEB journal : official publication of the Federation of American Societies for Experimental Biology 2021;35:e21722.

45. DeVera C, Dixon J, Chrenek MA, et al. The Circadian Clock in the Retinal Pigment Epithelium Controls the Diurnal Rhythm of Phagocytic Activity. International journal of molecular sciences 2022;23.

46. Vargas JA, Finnemann SC. Differences in Diurnal Rhythm of Rod Outer Segment Renewal between 129T2/SvEmsJ and C57BL/6J Mice. International journal of molecular sciences 2022;23.

47. LaVail MM. Rod outer segment disk shedding in rat retina: relationship to cyclic lighting. Science (New York, NY) 1976;194:1071–1074.

48. Possidente B, Hegmann JP, Carlson L, Elder B. Pigment mutations associated with altered circadian rhythms in mice. Physiology & behavior 1982;28:389–392.

49. Lynch CJ. The so-called Swiss mouse. Laboratory animal care 1969;19:214–220.

50. Schwartz WJ, Zimmerman P. Circadian timekeeping in BALB/c and C57BL/6 inbred mouse strains. The Journal of neuroscience : the official journal of the Society for Neuroscience 1990;10:3685–3694.

51. Meek KM, Knupp C. Corneal structure and transparency. Progress in retinal and eye research 2015;49:1–16.

